# Independent spatiotemporal effects of spatial attention and background clutter on human object location representations

**DOI:** 10.1101/2022.05.02.490141

**Authors:** Monika Graumann, Lara A. Wallenwein, Radoslaw M. Cichy

## Abstract

Spatial attention helps us to efficiently localize objects in cluttered environments. However, the processing stage at which spatial attention modulates object location representations remains unclear. Here we investigated this question identifying processing stages in time and space in an EEG and fMRI experiment respectively. As both object location representations and attentional effects have been shown to depend on the background on which objects appear, we included object background as an experimental factor. During the experiments, human participants viewed images of objects appearing in different locations on blank or cluttered backgrounds while either performing a task on fixation or on the periphery to direct their covert spatial attention away or towards the objects. We used multivariate classification to assess object location information. Consistent across the EEG and fMRI experiment, we show that spatial attention modulated location representations during late processing stages (>150ms, in middle and high ventral visual stream areas) independent of background condition. Our results clarify the processing stage at which attention modulates object location representations in the ventral visual stream and show that attentional modulation is a cognitive process separate from recurrent processes related to the processing of objects on cluttered backgrounds.

## 2 Introduction

Spatial attention helps us to focus visual processing on the relevant portions of the visual field while ignoring its irrelevant portions (Desimone and Duncan, 1995). For example, spatial attention helps during navigation to determine where in visual space objects are located, allowing us to avoid obstacles and to reach desired targets better.

While the importance of spatial attention is widely acknowledged, its neural basis remains incompletely understood. An important open question is, at which stage of the visual processing hierarchy attention modulates object location representations. Previous research yielded contradictory results. Considering the temporal emergence of attentional effects, some studies found attentional modulation early (Mangun, 1995; Hillyard et al., 1998a, 1998b; Luck et al., 2000) within a time window that corresponds to the initial bottom-up response within the first 150 ms (Lamme and Roelfsema, 2000; VanRullen and Thorpe, 2001; Fahrenfort et al., 2007; Camprodon et al., 2010; Koivisto et al., 2011) while others found such effects only later (Wyatte et al., 2014; Groen et al., 2016; Kaiser et al., 2016; Battistoni et al., 2020). Similarly, considering the locus in the visual processing hierarchy some studies found attentional modulation already in V1 (Roelfsema et al., 1998; Martínez et al., 2001; Noesselt et al., 2002; Khayat et al., 2006; Lakatos et al., 2008; Briggs et al., 2013; Herrero et al., 2013) while others found such effects only or predominantly in higher-level brain regions (Buffalo et al., 2010; Peelen and Kastner, 2011; Kay et al., 2015).

The contradiction might be resolved when considering together the processing stage at which object location representations emerge, the object’s viewing conditions and attentional modulation. Recent research has shown that viewing conditions influence the processing stage at which object location representations emerge. For example, object location representations emerge early for objects on blank and late on cluttered backgrounds (Hong et al., 2016; Graumann et al., 2022). Further, the surroundings of an object modulates the employment of spatial attention: spatial attention is more relevant for the localization of objects in clutter than in isolation (Treisman and Gelade, 1980; Wolfe, 1994).

Here we set out to untangle the complex link between the processing stage at which object location representations emerge, its viewing conditions, and the effect of attentional modulation.

Our hypotheses are as follows. We set the stage by hypothesizing based on recent findings that the processing stage at which object location representations emerge depends on the object’s viewing conditions in particular its background (Graumann et al., 2022) independent of spatial attention. This replication hypothesis was termed HReplication (abbreviated HR; Fig. 1A,C).

**Figure 1.**
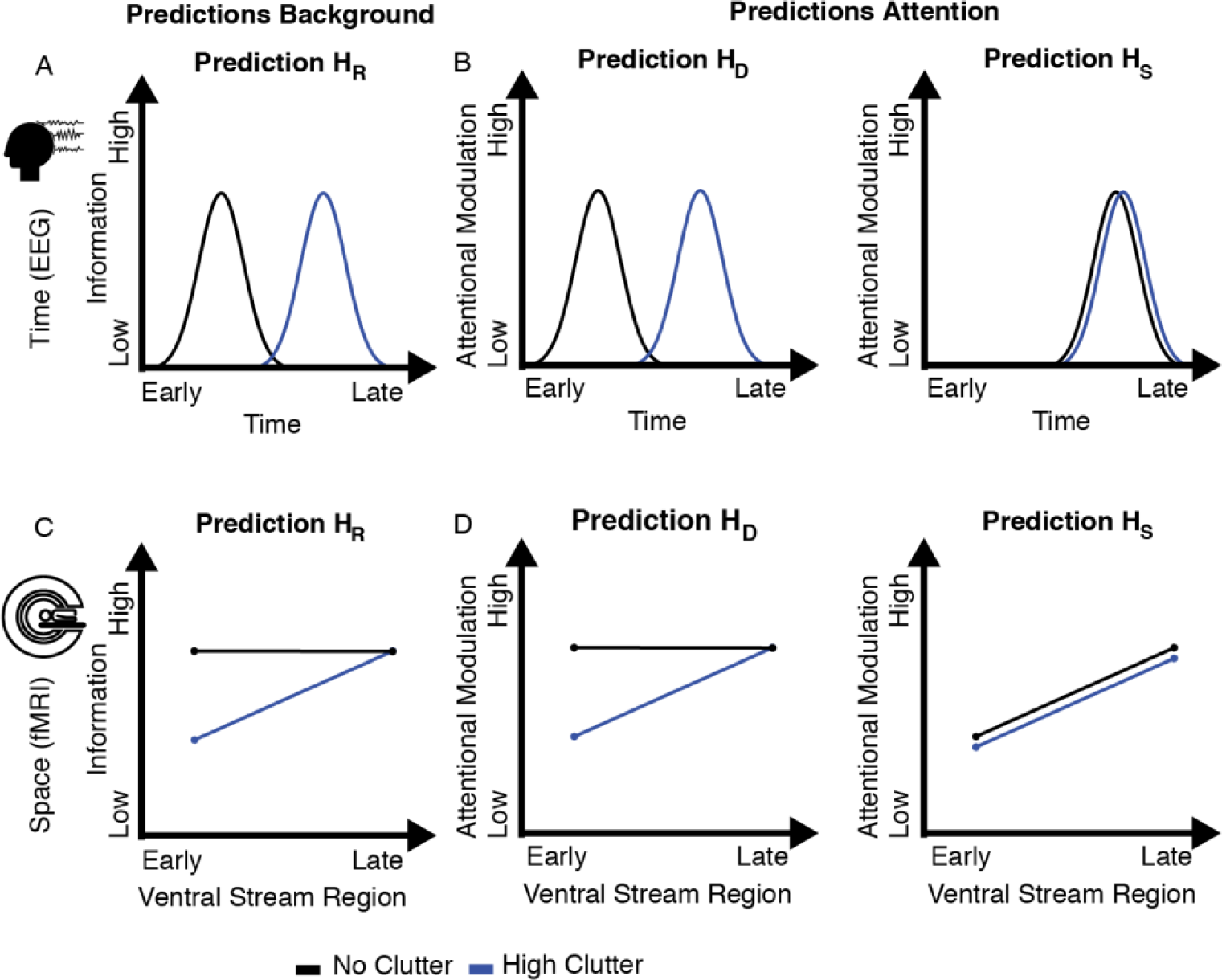
Experimental predictions based on hypotheses. A, Predictions for the effect of background on location information in the EEG experiment. HR predicts a delay in time for location information with high clutter compared to no clutter. **B,** Predictions for the effect of attention on location information in the EEG experiment. Predictions are based on HR in A. HD predicts that the time point when attentional modulation is highest depends on background: attentional modulation is highest at time points when location information is highest, depending on background condition. HS predicts that attentional modulation is always highest during late processing stages, independent of background condition. **C,** Predictions for the effect of background on location information in the fMRI experiment. HR predicts an increase along the ventral stream for location information with high clutter compared to no clutter. **D**, Predictions for the effect of attention on location information in the fMRI experiment. Predictions are based on HR in C. HD predicts that the region where attentional modulation is highest depends on background: attentional modulation is highest in regions where location information is highest, depending on background condition. HS predicts that attentional modulation always increases along the ventral stream, independent of background condition.

On this basis we then theorize how an objects background impacts when (in time with respect to stimulus onset) and where (in the cortical processing hierarchy) attention modulates location representations. We propose two alternative hypotheses.

The first hypothesis is that attention and background interact: attention dynamically modulates location representations at the processing stage at which they first emerge, resulting in an interaction between background and attention (HDynamic, abbreviated HD; Fig. 1B,D). The alternative hypothesis is that attention modulates location representations statically and always during a late processing stage (Wyatte et al., 2014; Kay et al., 2015; Groen et al., 2016; Kaiser et al., 2016; Battistoni et al., 2020), independent of the background (HStatic, abbreviated HS; Fig. 1B,D).

We investigated these hypotheses in an integrated research project consisting of an EEG and fMRI experiment in combination with multivariate pattern analysis methods. We manipulated background by presenting objects on backgrounds of different clutter levels, and attention by task instruction that attracted or diverted spatial attention from an object’s location.

To anticipate, we first confirmed HR, i.e., object location representations emerged later in time and space when the object appeared on a cluttered background than on a blank background, independent of attention. We then found strong empirical support for HS. That is, attention modulates object location representations late in both time and space, independent of background.

## 3 Results

For both the EEG and fMRI experiments the strategy to determine when and where attention modulates location representations was equivalent: first we sought to establish HR, i.e., that location representations of objects emerge at a later processing stage when objects are presented on cluttered backgrounds compared to blank backgrounds, independent of attention. On this basis we then arbitrated between HD and HS, i.e., whether attention dynamically modulates object location representations at different processing stages depending on background, or whether it statically modulated object location representations always at a late processing stage.

The difference between the EEG and the fMRI analyses lies in the way that the processing stages are determined: EEG determines the temporal delay with respect to image onset (Fig. 1A,B) and fMRI determines the region in the ventral visual stream (Fig. 1C,D) in which experimental effects emerge.

In the following we give the specifics of the EEG and fMRI experiments, the precise predictions, and the results. We begin with the EEG experiment determining the timing of attentional modulation, followed by the fMRI experiment determining where in the visual processing hierarchy the attentional modulation occurs.

### 3.1 Attentional modulation of object location representations in time

For the EEG experiment we used an experimental design with fully crossed experimental factors background and attention with two levels per factor (2 background condition levels × 2 attention condition levels, Fig. 2A).

**Figure 2.**
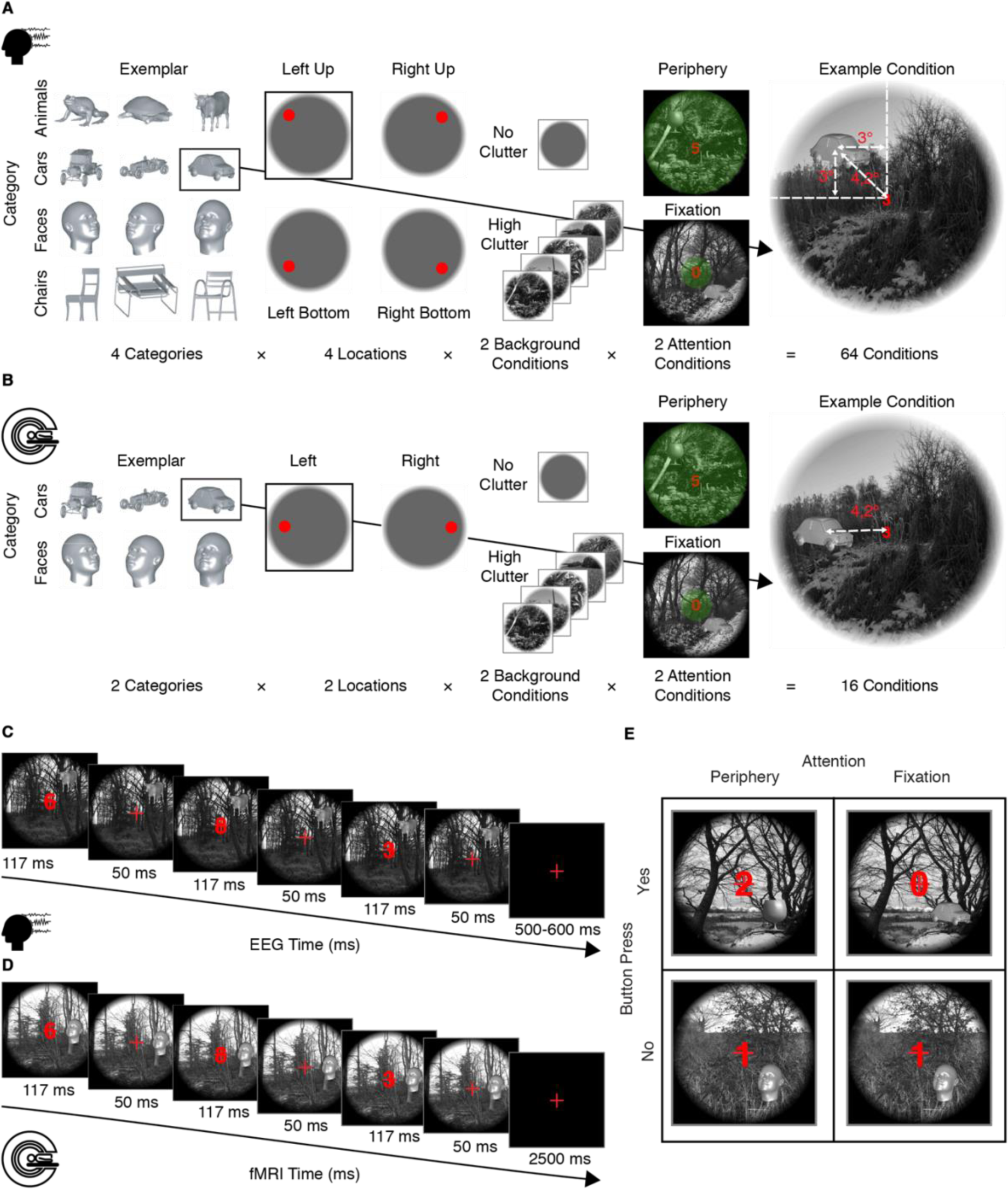
Experimental design and tasks. A, Experimental design in the EEG experiment. We used a fully crossed design with factors: object category, location, background and attention. Green translucent circles represent attentional width. **B**, Experimental design in fMRI experiment. The design was equivalent to the EEG experiment, except that the factors category and location had two levels. **C,** Trial timing and example condition in EEG experiment. **D,** Trial timing and example condition in fMRI experiment. **E,** Tasks. In the peripheral attention condition (left) participants responded with button press when a glass appeared in the periphery, while fixating their gaze on the central cross. Digits presented on fixation were task-irrelevant. In the fixation attention condition (right) participants responded with button press when the digit 0 appeared on fixation, while fixating on the central cross. Objects in the periphery were irrelevant in this task. Visual stimulation was the same in both tasks on regular trials (see bottom row ‘Button press: no’).

In detail, participants saw objects from four different categories, each presented in four locations (Fig. 2A). The two background conditions were no and high clutter. Each object in each location was presented in both background conditions. Each combination of object category, location and background was then also crossed with the two levels of attention conditions: the peripheral and fixation attention conditions (Fig. 2A). Attention conditions were solely defined by the task that participants performed, while visual stimulation was identical (Fig. 2E). In the peripheral attention condition, participants directed their covert spatial attention to the periphery and responded to a catch object (glass) with button press (Fig. 2A,E). In the fixation attention condition, participants performed a demanding task on fixation to remove their spatial attention from the objects in the periphery (Fig. 2A,E).

In total, this 2 × 2 experimental design resulted in 4 factor combinations. We performed a time-resolved and pair-wise classification analysis of location across category within each of these four factor combinations separately (Fig. 3A,B). This meant training a classifier to distinguish between millisecond-specific EEG pattern vectors associated with two locations and testing on a held-out testing data set associated with the same two locations. We performed the classification across object category, that is training on data associated with locations from one object category and testing on data from another category (Fig. 3A). This ensured that location classification results were not confounded with category information and allowed us to draw conclusion about location representations independent of object category representations.

**Figure 3.**
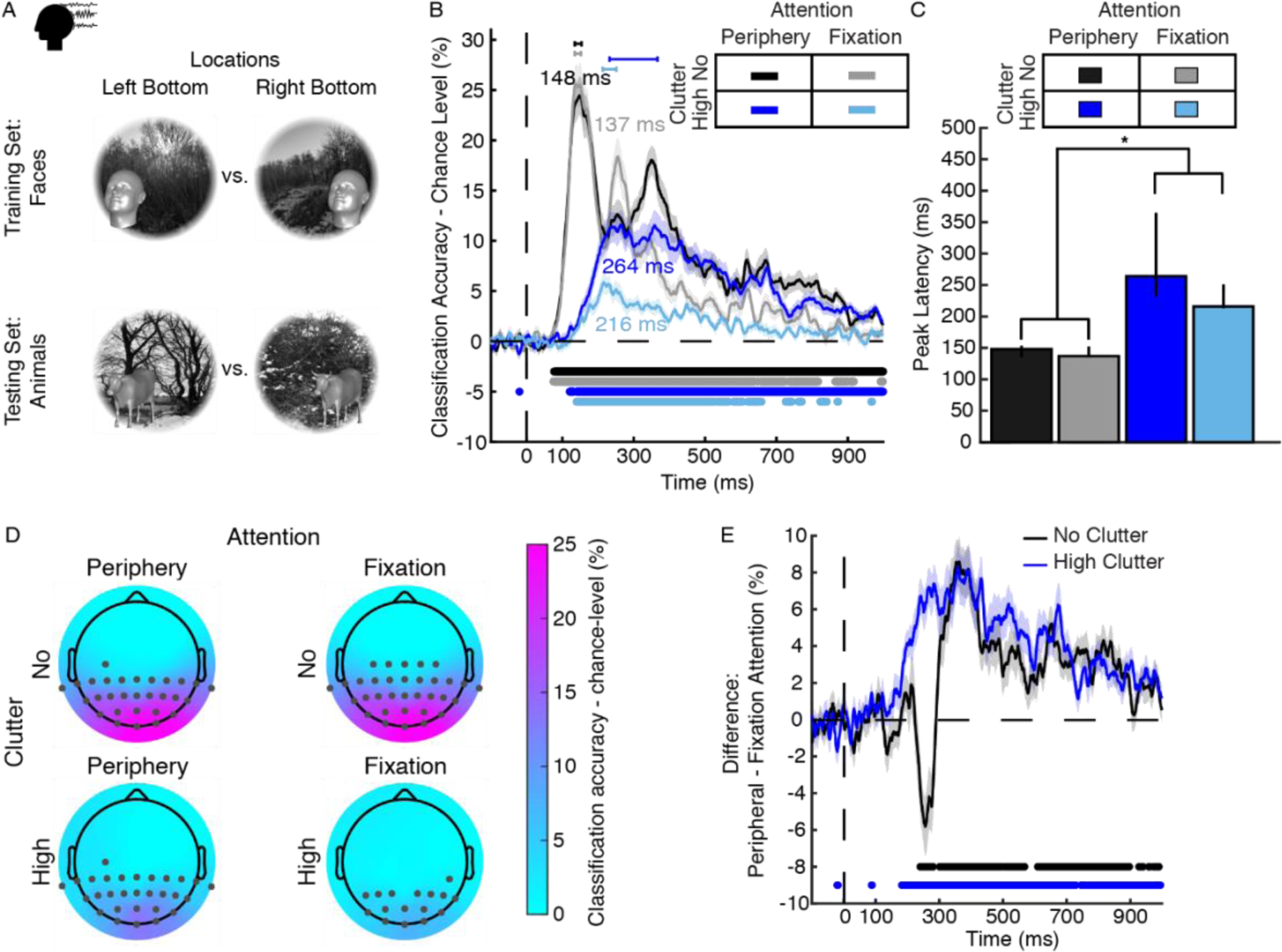
Classification schemes and results of EEG location classification. A, Scheme for the classification of object location across categories within any background and attention condition. We trained a support vector machine (SVM) to distinguish brain activation patterns evoked by objects of a particular category presented at two locations (here: faces bottom left and right) and tested the SVM on activation patterns evoked by objects of another category (here: animals) presented at the same locations. Objects are enlarged here for display purposes. In the experiment objects did not extend across quadrants. **B,** Results of time-resolved location across category classification from EEG data. Results are color-coded by background and attention condition, with significant time points indicated by lines below curves (*N*=26, *P*<0.05, FDR-corrected), 95% confidence intervals of peak latencies are indicated by lines above curves. Shaded areas around curves indicate SEM. **C,** Comparison of peak latencies of curves in B. Error bars represent 95% CIs. Stars indicate significant peak latency differences (*N*=26, bootstrap test with 10,000 bootstraps). **D,** Results of the location across category classification searchlight in EEG channel space at peak latencies (as shown in B) in each condition. Significant electrodes are indicated by grey dots (*N*=26, two-tailed Wilcoxon signed-rank test, *P*<0.05, FDR-corrected across electrodes and time points). **E,** Difference curves resulting from subtracting the time courses of the foveal from the peripheral attention condition in each background condition. Conventions as in B.

#### 3.1.1 The temporal dynamics of object location representations with blank and cluttered backgrounds

To lay the basis for later analyses on attentional modulation, we first tested HR, i.e., that location representations of objects with clutter emerge later than on blank backgrounds, independent of attention. For this we determined and compared the latencies of the classification peaks in the EEG time courses of both background conditions, assuming that the peaks represent the time points at which representations become most differentiable (DiCarlo and Cox, 2007). Our prediction was that location information would peak later in the high than in the no clutter condition, because dissecting objects from the background requires additional grouping and segmentation operations implemented in recurrent processing and thus increasing processing time (Groen et al., 2018; Seijdel et al., 2020, 2021; Graumann et al., 2022).

The results of the time-resolved location classification are shown in Fig. 3B. We read out location information from the EEG signal in all background and attention conditions above chance level (*N*=26, two-tailed Wilcoxon signed-rank test, *P*<0.05, FDR-corrected).

Focusing on peak latencies (95% confidence intervals reported in brackets, *N*=26, 10,000 bootstrap samples), we observed that time courses in the no and high clutter conditions peaked at different times. In the no clutter condition, location information peaked early, regardless of attention condition (Fig. 3B; peak latency peripheral condition: 148 ms (135–153.5 ms); peak latency fixation condition: 137 ms (135–152 ms)). With high clutter, location information peaked later in both attention conditions (Fig. 3B; peripheral condition: 264 ms (232–365 ms); fixation condition: 216 ms (213–251 ms)). To test whether the peak latencies across background conditions were significantly different, we bootstrapped the peak-to-peak latency differences between pairs of no and high clutter condition peaks (Fig. 3C, 95% confidence intervals in brackets, *N*=26, bootstrap test, 10,000 bootstraps, FDR-corrected). This was done both within and across attention conditions. Overall, the results clearly and consistently support HR. Location information peaked significantly earlier in the no compared to the high clutter conditions independent of attention condition: Within attention condition, the peak-to-peak latency difference between background conditions was 116 ms (83–223 ms; *P*<0.001) in the peripheral attention condition and 79 ms in the fixation attention condition (63–114 ms; *P*<0.001). Across attention conditions, the delays between background condition peaks were also significant (peripheral attention and no clutter condition vs. fixation attention and high clutter condition: 68 ms delay, 63–105 m; *P*<0.001; fixation attention and no clutter condition vs. peripheral attention and high clutter condition: 127 ms delay, 83–224 ms, *P*<0.001).

Additional analyses of the observed effects reproduced previously observed characteristics of object location representations (Graumann et al., 2022) and thus further supported HR. A searchlight analysis in EEG sensor space (Fig. 3D) localized the sources of the peaks to occipito-temporal electrodes (Fig. 3D), suggesting the locus of object location representations to be in occipital and temporal cortices. A supplementary time-generalization analysis (King and Dehaene, 2014) showed that location representations for objects on blank and cluttered background emerged within the same processing stage, but with a delay with cluttered backgrounds (Supplementary Fig. 1, Supplementary Methods 1).

Together, these results provide empirical evidence for HR.

#### 3.1.2 Late attentional modulation of location representations independent of background

Affirming HR formed the basis for arbitrating between our main hypotheses HD and HS. HD predicts that attentional modulation is highest when location information is highest: with no clutter, it predicts an early modulation in time of location representations and with high clutter it predicts a late modulation in time (Fig. 1B). HS states that spatial attention modulates location representations always late, after the end of the bottom-up response at ∼100-150 ms (Lamme and Roelfsema, 2000; VanRullen and Thorpe, 2001; Fahrenfort et al., 2007; Camprodon et al., 2010; Koivisto et al., 2011). Thus, HD predicts an interaction between attention and background and HS predicts that they are independent.

To assess HS and HD we determined the time course of attentional modulation in both background conditions. Attentional modulation was defined as an enhancement of representations (Desimone and Duncan, 1995; Reynolds and Chelazzi, 2004; Briggs et al., 2013). To quantify attentional modulation, we subtracted classification accuracies in the fixation attention condition from the peripheral attention condition, within each background condition. Since visual stimulation was identical across attention conditions, we attributed differences between them to attentional modulation.

Fig. 3E shows the result of this analysis. We found attentional modulation of location representations in both background conditions in a late time window, providing clear evidence for HS. In detail, we observed a significant positive difference in the no clutter condition starting from ∼300 ms, reflecting attentional modulation (Fig. 3E; *N*=26, two-tailed Wilcoxon signed-rank test, *P*<0.05, FDR-corrected). In the high clutter condition, we found evidence for attentional modulation starting from 182 ms (Fig. 3E; *N*=26, two-tailed Wilcoxon signed-rank test, *P*<0.05, FDR-corrected), as reflected in a significant positive difference that lasted until the end of the time window.

Together, these results show that attention modulates object location representations in a late time window after the bottom-up response, independent of background. This constitutes strong evidence for HS.

#### 3.1.3 Dissecting transient and persistent components of attentional modulation

While clearly supporting HS, the results hitherto do not yet characterize the temporal dynamics underlying attentional modulation of location representations. Typically during visual perception, time-resolved multivariate results reflect a conglomerate of both rapidly changing transient information flow as well as persistent activity which maintains certain types of information over long stretches of time (Cichy et al., 2014; King and Dehaene, 2014).

Thus, here we investigated whether attention and background modulate persistent, transient or both aspects of location representations. For this we conducted temporal generalization analysis (King and Dehaene, 2014). This resulted in two-dimensional time generalization matrices, indexed in both dimensions in time indicating similarities of object location representation across time. While transient representations are reflected as high information on the diagonal of such matrices, persistent representations are found off-diagonal.

As previously, we classified location representation within background and attention condition, resulting in 4 time-generalization matrices (Fig. 4A,B,D,E), corresponding to the 4 classification time courses above (Fig. 3B). We first present the single results ordered by background condition, before quantifying attentional modulation.

**Figure 4.**
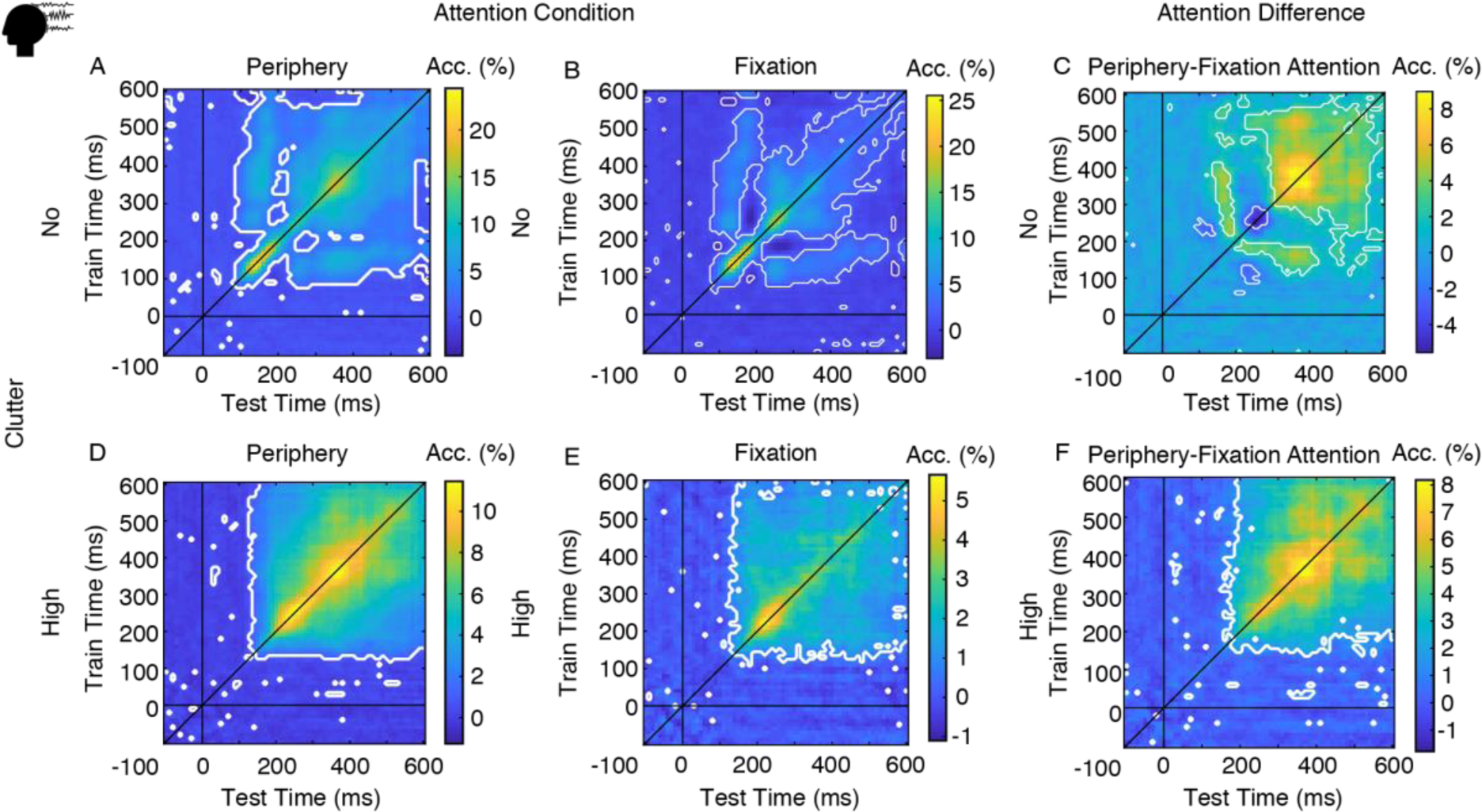
EEG results of time-generalization analyses within each background and attention condition. Rows represent background and columns represent attention conditions. **A,** Location classification across categories and time points in the no clutter & peripheral attention condition. Horizontal and vertical black lines indicate stimulus onset, oblique black line highlights the diagonal. White outlines indicate significant time points (*N*=26, two-tailed Wilcoxon signed-rank test, *P*<0.05, FDR-corrected). **B,** Location classification across categories and time points in the no clutter & fixation attention condition. **C,** Difference matrix resulting from subtracting the matrices representing fixation (B) from peripheral attention (A) in the no clutter condition. Plot conventions as in A. **D,** Location classification across categories and time points in the high clutter & peripheral attention condition. **E,** Location classification across categories and time points in the high clutter & fixation attention condition. **F,** Difference matrix resulting from subtracting the matrices representing fixation (E) from peripheral attention (D) in the high clutter condition.

In the no clutter condition, we found similar results in both attention conditions (Fig. 4A,B): location information peaked early at ∼100 ms on the diagonal, representing transient information flow (*N*=27, *P*<0.05, two-tailed Wilcoxon signed-rank test, FDR-corrected). Starting from ∼250 ms, location information generalized more broadly across time points, indicating persistent information. In the high clutter condition (Fig. 4D,E) information generalized broadly across time points starting from ∼140 ms in both attention conditions, (*N*=27, *P*<0.05, two-tailed Wilcoxon signed-rank test, FDR-corrected), indicating persistent information. Transient information peaked on the diagonal starting from ∼240 ms.

We quantified attentional modulation as above (Fig. 3E) by comparing the classification results for the two attention conditions, subtracting the results of the fixation attention condition from the peripheral attention condition.

We found that spatial attention modulated both transient and persistent representations in late time windows, independent of background. In the no clutter condition, attention modulated the persistent clusters from ∼230 ms and both transient and persistent information from ∼300 ms (Fig. 4C; *N*=27, *P*<0.05, two-tailed Wilcoxon signed-rank test, FDR-corrected). In the high clutter condition, spatial attention modulated location representations across the entire time window starting from ∼180 ms (Fig. 4F; *N*=27, *P*<0.05, two-tailed Wilcoxon signed-rank test, FDR-corrected).

In sum, we found that attention modulates both transient and persistent representations of object location representations in late time beyond 150 ms.

### 3.2 Clutter and attention independently affect location representations along the ventral visual stream

We proceed to investigate which visual processing stages are modulated by background and attention in an fMRI experiment, determining processing stages by localizing and assessing cortical regions of the ventral visual stream (Fig. 1C,D). In this context HR predicts that location representations of objects emerge in higher regions along the ventral stream when objects are presented on cluttered backgrounds compared to blank backgrounds (Graumann et al., 2022; Fig. 1C). HD predicts that attentional modulation is high where location information is high (Fig. 1D): with no clutter, attention modulates location representations throughout the ventral stream and with high clutter attention modulates location representations in mid- or high-level visual areas. HS instead predicts that attentional modulation is high in mid- and high-level visual areas only, independent of background.

The design of the fMRI experiment was equivalent to the design in the EEG experiment with a reduced number of levels for the factors category and location. This adaptation was made to accommodate the longer trial duration required for our fMRI event-related design (Fig. 2D) while maintaining a feasible session duration. We presented objects from two categories (faces, cars) in two locations (left and right horizontally from fixation; Fig. 2B) instead of four categories and locations. To characterize the effects of background and spatial attention on location representations in visual cortex, we defined regions-of-interest (ROIs) along the ventral stream, since HR predicted effects to emerge there based on previous studies (Hong et al., 2016; Graumann et al., 2022). We additionally included ROIs in the dorsal stream, since it has been implicated in visuospatial (Ungerleider and Haxby, 1994; Milner and Goodale, 2006; Kravitz et al., 2011; Groen et al., 2022) and attentional processing (Silver et al., 2005; Szczepanski et al., 2010; Sprague and Serences, 2013).

We classified location across category using an analogous classification scheme as in the EEG experiment. We trained a classifier on fMRI patterns associated with two locations of one object category and subsequently cross-validated the classifier on new testing data associated with the same locations of a new object category. This classification was performed in each ROI separately, for each level of the background condition (no clutter, high clutter) and for each level of attention conditions (periphery, fixation) separately, resulting in four classification accuracies per ROI and subject. We included three ROIs in early visual cortex (V1, V2, V3), two ROIs in the ventral stream (V4, LOC) and four ROIs in the dorsal stream (IPS0, IPS1, IPS2, SPL).

We tested HR, HS and HD in 2 × 2 repeated-measures ANOVAs (*N*=20, FDR-correction for multiple comparisons) with factors background (no clutter, high clutter) and attention (peripheral, fixation) in all ROIs of the ventral and dorsal visual streams, focusing on the ventral visual stream first.

Hypothesis HR predicted a main effect of background in early visual areas, but not in high-level visual areas of the ventral visual stream. Consistent with the predictions of HR, we found significant main effects of background in early visual areas V1 and V2 (Fig. 5A,B; V1: *F*(1,19)=9.88, *P*=0.005, partial *η^2^*=0.34; V2: *F*(1,19)=11.56, *P*=0.003, partial *η^2^*=0.3), but not in mid- and high-level visual areas V3 and LOC (Table 1), except for V4, which also showed a main effect of background (*F*(1,19)=16.64, *P*<0.001, partial *η^2^*=0.47).

**Figure 5.**
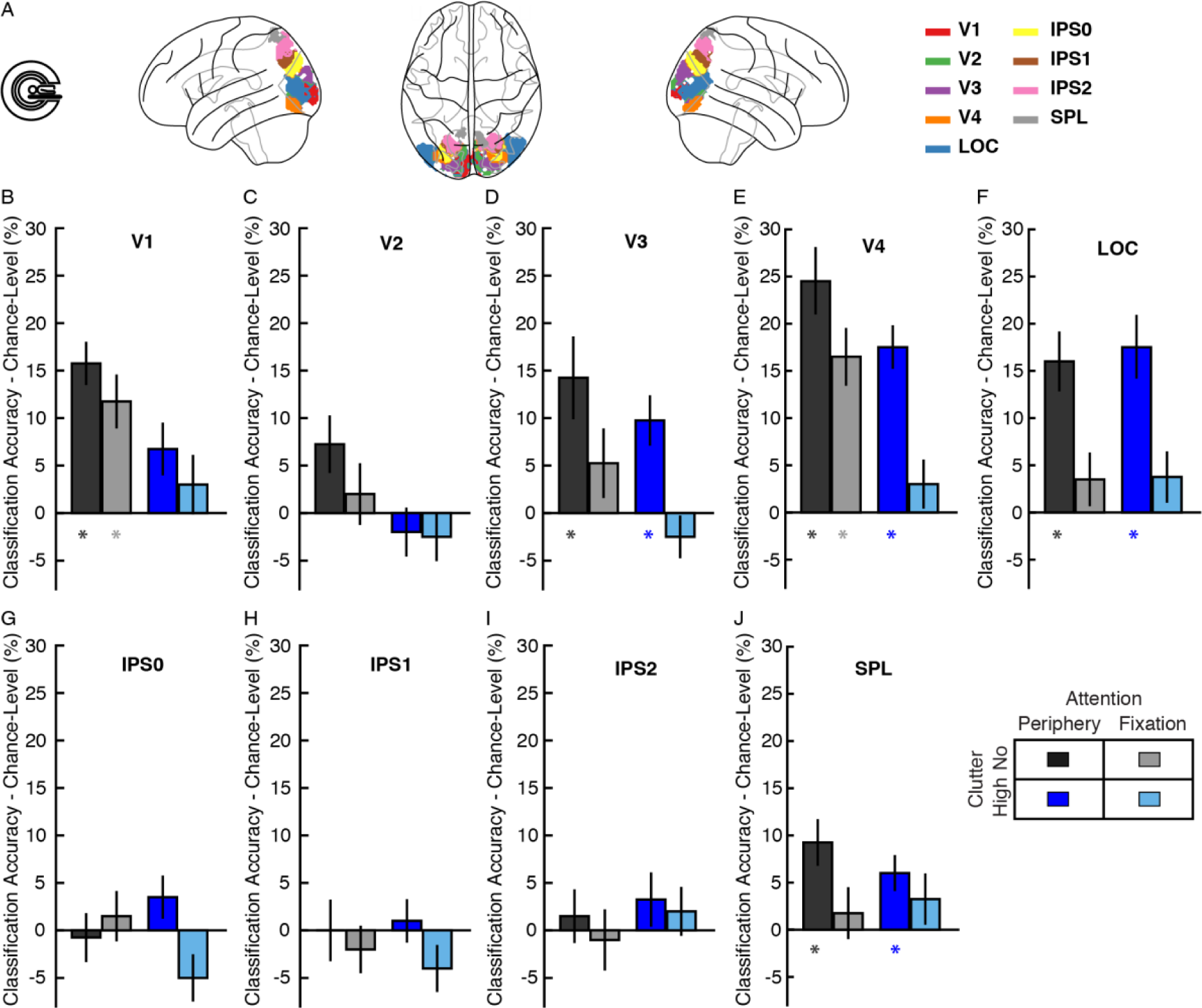
Location across category classification results in the four conditions in early (V1, V2, V3), ventral (V4, LOC) and dorsal (IPS0-2, SPL) visual ROIs. Stars below bars indicate significant above-chance classification (*N*=20, two-tailed Wilcoxon signed-rank test, *P*<0.05 false discovery rate (FDR) corrected). Error bars represent standard error of the mean (SEM). **A,** ROIs on cortical surface. **B,** V1. **C,** V2. **D,** V3. **E,** V4. **F,** LOC. **G,** IPS0. **H,** IPS1. **I,** IPS2. **J,** SPL.

**Table 1.**
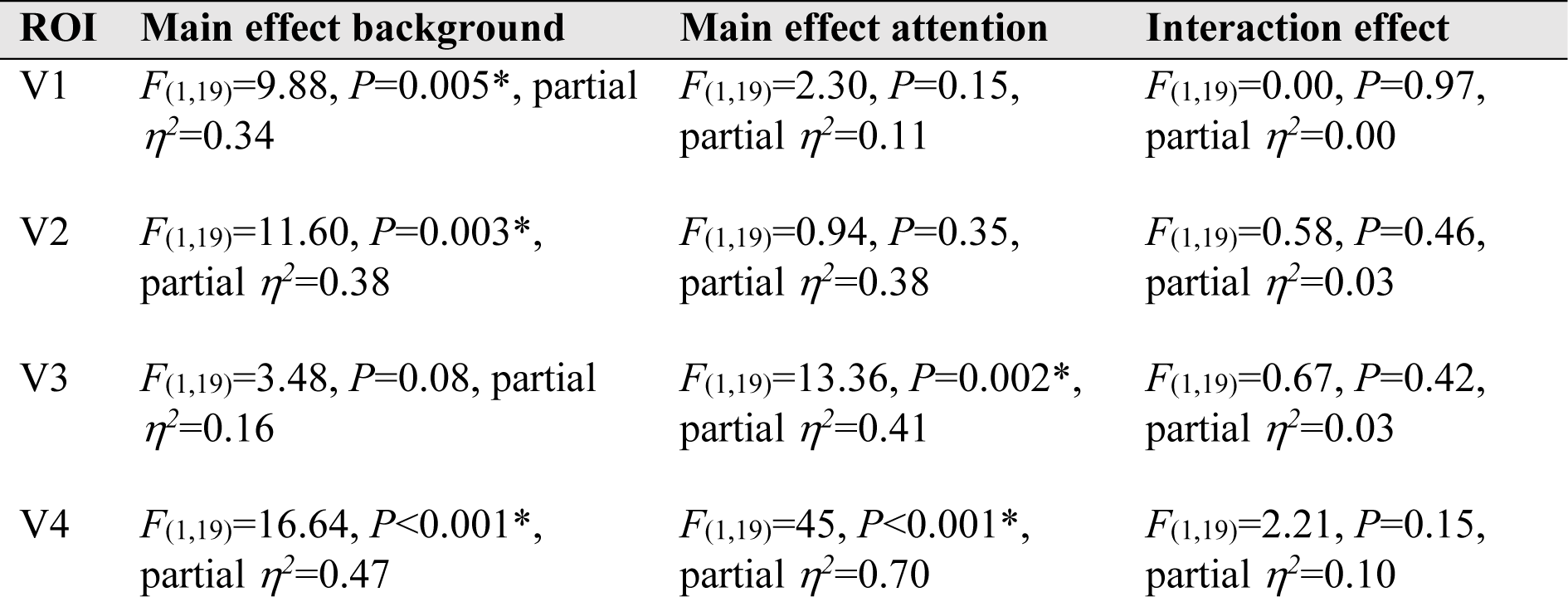
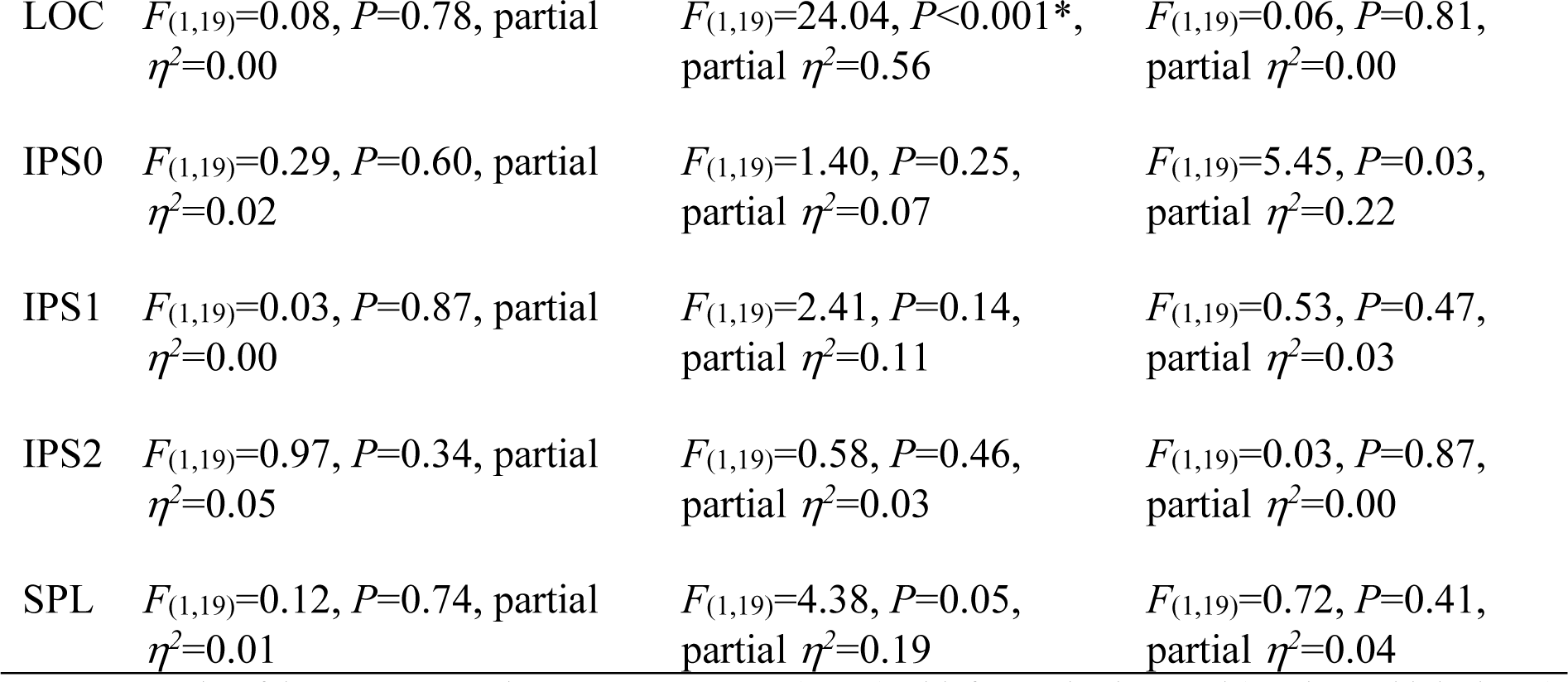
Results of the 2 × 2 repeated-measures ANOVA (*N*=20) with factors background (no clutter, high clutter) and attention (peripheral, fixation), analyzing location classification accuracies in 9 ROIs that were included in the analyses. Asterisks behind *P*-values indicate significance with FDR correction across number of comparisons (for 9 ROIs).

On this basis arbitrating between HS and HD we found clear evidence for HS. Location information in mid- and high-level ventral visual areas V3, V4 and LOC all showed a significant main effect of attention (Fig. 5C,D,E; V3: *F*(1,19)=13.36, *P*=0.002, partial *η^2^*=0.41; V4: *F*(1,19)=45, *P*<0.001, partial *η^2^*=0.70; LOC: *F*(1,19)=24.04, *P*<0.001, partial *η^2^*=0.56), but no significant interaction between background and attention as would have been predicted by HD.

Equivalent testing in the dorsal visual stream revealed no significant main or interaction effect in any regions along the dorsal stream (Fig. 5F,G,H,I; Table 1), consistent with the observation that object location representations emerge rather along the ventral than the dorsal stream (Hong et al., 2016; Graumann et al., 2022).

In sum, the results of the fMRI experiment concur with the results of the EEG experiment in providing evidence for HR and HS. Object location representations emerge gradually along the processing hierarchy of the ventral visual stream, and attention modulates object location representations in mid- and high-level ventral areas independent of the object’s background.

## 4 Discussion

Using EEG and fMRI we investigated at which stage of the visual processing hierarchy attention modulates object location representations. Our results converge across the two experiments and imaging modalities into a common view. We reproduced the recent observation that object location representations emerge at later processing stages when presented on cluttered than on blank backgrounds (HR) and showed that this holds independent of attention. On this basis we examined the effect of attention on object location representations, finding that attention modulated location representations statically during late stages of visual processing in cortical time and space, independent of the object’s background (HS).

### 4.1 Disentangling the influences of background and attention on the temporal dynamics of location representations

Recent research has revealed that object location representations emerge later in the ventral visual processing hierarchy when objects appear on cluttered rather than on blank backgrounds (Hong et al., 2016; Graumann et al., 2022). However, it remained unclear to which degree this effect relied on or was influenced by attention or not. Previous research has highlighted attention as important for object perception under cluttered conditions (Treisman and Gelade, 1980; Wolfe, 1994; Reddy and Kanwisher, 2007; Lee and Maunsell, 2010). Further, both temporal delays observed for object perception and attention have been related to recurrent processes (Tang et al., 2014; Kar et al., 2019; Rajaei et al., 2019; van Bergen and Kriegeskorte, 2020), suggesting shared neural mechanisms.

Here, we clarify the relationship and find a dissociation: while cluttered viewing conditions delay processing (see also time-generalization analysis in Supplementary Fig. 1, Supplementary Methods 1), spatial attention in contrast increases information without changing its timing. These results suggest that background clutter and attention have differential effects on object location processing. Background clutter, like other factors that increase image complexity, triggers local recurrent processes that can be measured in delayed responses (Tang et al., 2014, 2018; Groen et al., 2018; Kar et al., 2019; Rajaei et al., 2019; Seijdel et al., 2021; Graumann et al., 2022). In contrast, spatial attention triggers modulation of neural responses that can be measured as enhancement of response magnitude (Desimone and Duncan, 1995; Reynolds and Chelazzi, 2004; Briggs et al., 2013).

### 4.2 Attention modulates location representations later than the initial bottom-up response

The EEG results revealed attentional modulation of location representations in a late time window beyond the first 150 ms of the bottom-up response independent of the object’s background. In-depth investigation further revealed that both transient and persistent neural components were modulated. This directly supports the hypothesis Hs that the processing stage of attentional modulation is static and refutes the hypothesis HD that the processing stage at which attention modulates location representations changes dynamically.

Our results are seemingly at odds with earlier studies finding attentional modulation before 150 ms in the P1 (Hillyard et al., 1998b; Luck et al., 2000; Itthipuripat et al., 2019) and the N1 (Mangun, 1995; Hillyard et al., 1998a; Itthipuripat et al., 2019) component. How is this discrepancy to be explained? We believe that the viewing conditions and the choice of the stimulus are relevant. Above mentioned ERP studies employed simple artificial stimulation conditions which might elicit attentional modulation already early. However, later studies using naturalistic stimuli, comparable to the ones used here, did not find early attentional modulation (VanRullen and Thorpe, 2001; Groen et al., 2016; Kaiser et al., 2016; Battistoni et al., 2020). Together this questions the degree to which previously observed effects of early attentional modulation generalize to more complex stimuli and naturalistic viewing conditions encountered in the real world.

Another contributing factor to the discrepancy could be that attentional enhancement of early neural responses is stronger when the visual task is more difficult or when visual processing is overloaded (Spitzer et al., 1988; Lavie, 1995; Luck et al., 2000; Boudreau et al., 2006; Chen et al., 2008) which might have been the case in earlier ERP studies e.g. by presenting stimuli in faster sequences (Hillyard et al., 1998b). In contrast in our experiment, stimuli in the no clutter condition were highly salient and presented long enough to be clearly visible. Future research comparing attentional modulation of artificial vs. real-world stimuli with different levels of task difficulty are needed to resolve this issue.

### 4.3 Attentional modulation in mid- and high-level ventral visual areas

Consistent with the EEG results indicating attentional modulation of later visual object processing stages in time, the fMRI experiment localized those modulations to mid- and high-level ventral visual areas. While our results do not exclude the existence of attentional modulation also in early visual cortex as observed previously (Roelfsema et al., 1998; Gandhi et al., 1999; Martínez et al., 2001; Noesselt et al., 2002; Khayat et al., 2006; Lakatos et al., 2008; Briggs et al., 2013; Herrero et al., 2013; Itthipuripat et al., 2019), they suggest that the modulation might be strongest and thus most likely to be detected at later stages of ventral visual cortex (Murray and Wojciulik, 2004; Buffalo et al., 2010; Peelen and Kastner, 2014; Kay et al., 2015). Our results add further evidence towards the view that attentional modulation begins in higher processing stages and is then relayed back to lower stages (Buffalo et al., 2010), which could be reflected as increasing attentional modulation along the ventral stream (Kay et al., 2015).

### 4.4 Location representations of objects on cluttered backgrounds in the ventral stream

The fMRI results reveal a double dissociation between the effects of clutter and attention on early and late ventral visual areas: early visual areas show an effect of background but not of attention, while the reverse is true for mid- and high-level visual areas. Put differently, we find that both robustness to clutter (Hong et al., 2016; Graumann et al., 2022) and attentional modulation increase along the ventral visual stream (Buffalo et al., 2010; Kay et al., 2015). We speculate that these phenomena depend on the common mechanistic and computational basis of receptive field size increases along the ventral visual stream. Attention increases population receptive field (pRF) size in higher-level ventral areas, thereby enhancing location sensitivity (Kay et al., 2015). Such pRF size increase might also simultaneously benefit object segmentation from cluttered backgrounds by encoding object location in global voxel patterns (Eurich and Schwegler, 1997; Kay et al., 2015). This benefit for object segmentation might in contrast not be present in early visual cortex where pRF size is small (Wandell and Winawer, 2015) and cells respond in a location unspecific way across all stimulated portions of the visual field to both objects and background clutter.

### 4.5 Limitations

We highlight two limitations of our experimental designs that are important for the correct interpretation of the results.

The first limitation is that in our experiment object locations and the content of the background are randomly paired and thus incongruent. In contrast, in the real world objects typically appear in locations congruent with the background scene. Attentional selection can exploit such relations between objects and backgrounds (Wolfe et al., 2011; Kaiser et al., 2019; Võ et al., 2019; Battistoni et al., 2020) on the basis of scene gist information (Oliva, 2005; Greene and Oliva, 2009). In our experiment this type of information cannot be exploited. Thus, when object locations and scene background are congruent, attentional modulation might be faster than revealed here. The flipside of the limitation is that our experimental design isolates the effect of clutter on visual processing and attentional modulation independent of congruency effects. To determine the effect of congruency of object location and background on visual processing, studies are needed that additionally investigate congruency as an experimental factor.

Another limitation is that we did not directly assess the behavioral effects of attentional modulation on localization performance. Spatial attention benefits object localization in cluttered displays (Treisman and Gelade, 1980; Wolfe, 1994; Wolfe et al., 2011) by increasing processing speed. Future studies may combine assessment with brain imaging to link the effect of attention for objects on cluttered backgrounds in brain and behavior.

### 4.6 Conclusion

In daily life, we use our spatial attention to help us focus on relevant portions of the visual field in cluttered environments (Wolfe et al., 2011). Our results clarify that attention modulates object location representations at late processing stages, using both spatial and temporal markers. Furthermore, they establish that attentional modulation is a cognitive process which is separate from recurrent processes which are engaged when objects appear in cluttered environments.

## 5 Materials and Methods

### 5.1 Participants in EEG and fMRI experiment

27 participants completed the EEG experiment. One participant was excluded because of technical problems, resulting in 26 participants (mean age 26.42 years, *SD*=4.12, 19 female) included in the final EEG study. 23 participants completed the fMRI experiment, out of which one also participated in the EEG experiment. Three participants were excluded because they did not complete the whole experiment, resulting in 20 participants (mean age 26.71 years, *SD*=4.48, 13 female) included in the final fMRI study.

All participants had no history of neurological disorders and normal or corrected-to-normal vision. Participants provided informed consent prior to the studies and participation was compensated with payment or course credit. The study was conducted in accordance with the Declaration of Helsinki and the ethics committee of the Department of Education and Psychology of the Freie Universität Berlin approved the study in advance.

### 5.2 Experimental design

#### 4.2.1 EEG experimental design

The experimental design in the EEG study comprised the four factors object category (animals, cars, faces, chairs, Fig. 2A left, with 3 exemplars per category), location (left up, left bottom, right bottom, right up, Fig. 2A left center), background (no and high clutter, Fig. 2A center) and attention (on periphery or on fixation, Fig. 2A right center). These four factors were fully crossed, to investigate them independently of each other. In total, this created 192 individual conditions (12 object exemplars × 4 locations × 2 background conditions × 2 attention conditions). For further analysis, data was collapsed across exemplars, so that data was analyzed at the level of category. Thus, the number of conditions for further analysis was 64 (4 categories × 4 locations × 2 background conditions × 2 attention conditions, Fig. 2A right).

#### 4.2.2 fMRI experimental design

The experimental factors in the fMRI experiment were the same as in the EEG experiment, but there were two instead of four levels for the factors category (cars, faces) and location (left, right; Fig. 2B). This resulted in 48 individual conditions (6 object exemplars × 2 locations × 2 background conditions × 2 attention conditions). For further analysis, data was likewise collapsed across exemplars, so that data was analyzed at the level of category. Thus, the number of conditions for further analysis was 16 (2 categories × 2 locations × 2 background conditions × 2 attention conditions).

### 5.3 Stimulus set generation

#### 4.3.1 Stimulus set generation: EEG experiment

The experimental design in the EEG study comprised 96 individual stimulus conditions shown in each attention condition, as detailed in the previous section. To create these stimuli, each exemplar was superimposed onto backgrounds with or without scene images in four locations. First, to position object exemplars onto the four image locations, we projected the 3D rendered objects onto to the four quadrants of the screen (Fig. 2A, left center). Rendered objects did not extend beyond a quadrant. Each object’s center was positioned 3 degrees from the vertical and 3 degrees from the horizontal central midline (i.e., 4.2 degrees diagonally from image center to fixation, Fig. 2A right), subtending 2.4 degrees (*SD*=0.4) in vertical and 2.2 degrees (*SD*=0.6) in horizontal extent.

Second, each exemplar in each location was superimposed onto a background with no and with high clutter (Fig. 2A, center; the backgrounds shown here are comparable to the original backgrounds used in the experiments). We chose the background conditions no and high clutter to compare visual stimuli with low and high image complexity, respectively (Groen et al., 2018). The no clutter condition was a uniform gray background. In the high clutter condition, we selected 60 natural scene images from the Places365 database (http://places2.csail.mit.edu/download.html) that did not contain objects of the categories included in our experimental design (i.e., no animals, cars, faces, chairs) and were highly cluttered (as defined by 10 independent subject ratings; for methods and results see Graumann et al., 2022). We converted the images to grayscale and superimposed a circular aperture of 15 degrees. Original backgrounds are not shown because of copyright reasons but are available here: https://osf.io/85sak/?view_only=db183dde8f4b406aaba5dfc0dd0ae67d.

From the set of 60 scene images, we selected 48 scene images to go with the 48 stimulus conditions within the high clutter condition (12 exemplars × 4 locations). To avoid systematic congruencies between objects and background images within the high clutter condition, stimulus conditions and backgrounds were randomly paired for each of the 20 runs into which the EEG experiment was divided (see below). Together with the 48 stimulus conditions in the no clutter condition, this resulted in 96 individual images per run. The 12 remaining scene images from the set of 60 were used to create catch trials.

#### 5.3.2 Stimulus set generation: fMRI experiment

Stimulus set generation for the fMRI experiment was equivalent to the EEG experiment, with the difference that objects were positioned on two instead of four image locations (Fig. 2B) 4.2° to the left or right of the image’s center. In the fMRI experiment, each background condition had 12 individual stimulus conditions (6 exemplars × 2 locations). In combination with the 12 stimulus conditions in the no clutter condition, this resulted in 24 individual images per run. The 12 remaining scene images from the set of 60 were used to create 24 catch trials (1 catch object × 12 scene images × 2 locations), which were randomly presented during the fMRI experiment.

### 5.4 Experimental procedures

#### 5.4.1 EEG main experiment

Each of the 26 participants completed one EEG recording session with 20 runs (run duration: 277 s). Overall, the EEG session lasted for 92 minutes. Participants performed attention tasks on separate runs. The EEG recording session consisted of 10 periphery attention runs and 10 fixation attention runs in randomized order. Within each attention condition, there were 96 individual stimulus conditions (12 exemplars × 4 locations × background conditions). Runs consisted of the presentation of regular trials and catch trials. In each run, there were 192 regular trials, representing the 96 stimulus condition images were presented twice. These trials formed the basis for further analysis. On regular trials, digits between 1 and 9 were overlaid for 117 ms each, followed by a 50 ms presentation of the image and fixation cross after each digit (Fig. 2C). In total, stimuli were presented for 0.5 s followed by 0.5 or 0.6 s of ISI (equally probable; Fig. 2C). Participants were asked to fixate their eyes on the central cross at all times.

On catch trials, a target was presented to which participants were asked to respond with button press (Fig. 2E). These trials were excluded from the analyses. Catch trials were presented on every 3^rd^ to 5^th^ trial (equally probable, in total 48 per run). Participants were instructed to respond with button press to catch trials and to blink their eyes to minimize eye blink contamination on subsequent trials. The ISI was 1s on catch trials to avoid contamination of movement and eye blink artefacts on subsequent trials.

In the periphery and the fixation attention condition different trials were task-relevant catch trials. In the periphery attention condition, catch trials were trials during which a target object (a glass) was presented (Fig. 2E). The target could be presented at any of the four locations and on any type of background. Digits on fixation were task-irrelevant in this attention condition. In the fixation attention condition, catch trials were trials during which the digit 0 appeared among any of the 3 digits that were presented on fixation during a single trial (Fig. 2E). The presented object in the periphery was task-irrelevant in this attention condition (Fig. 2E). The digit 0 never appeared on periphery attention runs and the glass never appeared on fixation attention runs.

#### 5.4.2 fMRI main experiment

Each of the 20 participants completed one fMRI recording session with 20 runs (run duration: 288 s). Overall, an fMRI recording in the main experiment lasted for 96 minutes. Each of the 24 images of the stimulus set was shown 3 times in random order without back-to-back repetitions in each run. On each trial, the image was presented for 0.5 s at the center of a black screen. The inter-stimulus-interval (ISI) was 2.5 s (Fig. 2D). Images were overlaid with a red central cross for fixation. Participants were instructed to fixate their eyes on this cross throughout the experiment. Every 3^rd^ to 5^th^ trial (equally probable, in total 18 per run) a catch trial was presented. The tasks in the attention conditions and the catch objects were identical to the EEG experiment (Fig. 2E). Catch trials were excluded from further analysis.

*fMRI localizer experiment.* Prior to the main fMRI experiment, participants completed a separate localizer run to define ROIs in early visual, dorsal and ventral visual stream. We presented images from three categories: faces, objects, and scrambled objects. Each image showed identical versions of the same object located left and right of fixation to stimulate the same retinotopic regions of visual cortex as the objects in the main experiment.

The localizer run lasted for 384 s, during which we presented 6 stimulation blocks. Each block was 16 s long with presentations of 20 different objects from one of the three categories (500 ms on, 300 ms off) block-wise. Each block included two one-back image repetitions to which participants had to respond to with a button press. The order of these blocks was first order counterbalanced: triplets of stimulation blocks were presented in random order and interspersed with blank background blocks.

### 5.5 EEG acquisition and preprocessing

To record EEG data, we used the EASYCAP 64-channel system with a Brainvision actiCHamp amplifier at a sampling rate of 1,000 Hz and with an online filter between 0.03 and 100 Hz. The signal was online re-referenced to FCz. Electrode placement followed the standard 10-10 system. Data was preprocessed offline with the EEGLAB toolbox version 14 (Delorme and Makeig, 2004). This comprised a low-pass filter with a 50 Hz cut-off, trial epoching in a peri-stimulus time window between -100 ms and 999 ms, and baseline-correction by subtracting the mean of the 100 ms prestimulus time window from the entire epoch. We used independent component analysis (ICA) to clean the data from ocular and muscular artefacts. To guide the visual inspection of components for removal we used SASICA (Chaumon et al., 2015). To identify horizontal eye movement components, we used external electrodes from the horizontal electrooculogram (HEOG). We detected blink artefact and vertical eye movements using the two frontal electrodes Fp1 and Fp2. On average, we removed 18 (*SD*=5) components per participant. We finally applied multivariate noise normalization on the preprocessed data to improve the signal-to-noise ratio and reliability of the data (Guggenmos et al., 2018).

### 5.6 Preprocessing and univariate fMRI analysis

*fMRI acquisition and preprocessing.* MRI data was recorded using a 12-channel head coil on a 3T Siemens Tim Trio Scanner (Siemens, Erlangen, Germany). The structural image was acquired with a T1-weighted sequence (MPRAGE; 1-mm^3^ voxel size). To acquire functional data for the main experiment and the localizer run, we ran a T2*-weighted gradient-echo planar sequence (TR=2, TE=30 ms, 70° flip angle, 3-mm^3^ voxel size, 37 slices, 20% gap, 192-mm field of view, 64 × 64 matrix size, interleaved acquisition) on the entire brain.

fMRI data was preprocessed using SPM8 (https://www._l.ion.ucl.ac.uk/spm/), involving realignment, coregistration and normalization to the structural MNI template brain. We smoothed functional data from the localizer run with an 8 mm FWHM Gaussian kernel, but the data from the main experiment were not smoothed.

*Univariate fMRI analysis.* We modelled the fMRI responses of the experimental conditions at the level of category. This was done for each run in the main experiment separately using a general linear model (GLM). We entered onsets and durations of stimulus presentations per category, pooling exemplars and repetitions. Thus, each GLM was estimated based on 9 trials (3 exemplars × 3 condition repetitions per run) and was convolved with the hemodynamic

response function (hrf). We further entered movement parameters into the GLM as nuisance regressors. This resulted in 8 beta maps per attention condition run (2 categories × 2 locations × 2 backgrounds). For each run, we converted GLM parameter estimates into *t*-values by contrasting each parameter estimate against the implicit baseline for each condition. This resulted for each participant and attention condition run separately in 8 (2 categories × 2 locations × background conditions) *t*-value maps per condition. In sum, this resulted in 8 *t*- value maps per 10 runs, per 2 attention conditions and per participant, which were later used in the classification analysis.

For the fMRI responses to the localizer experiment, we modelled the responses to objects, faces and scrambled objects by entering block onsets and durations as regressors of interest and movement parameters as nuisance regressors into the GLM and convolved them with the hrf. This resulted in three parameter estimates which we used to generate two contrasts that formed part of ROI definitions. The first contrast was defined as objects and scrambled objects > baseline and was used to localize activations in early, mid-level ventral and dorsal visual regions (V1, V2, V3, V4, IPS0, IPS1, IPS2, SPL). The second contrast was defined as objects and faces > scrambled objects and was used to localize activations in object-selective area LOC. Overall, this yielded two *t*-value maps for the localizer run for each participant.

*Definition of regions of interest.* To define ROIs, we first applied anatomical masks and then selected voxels using appropriate contrasts from the functional localizer run. In detail, we first defined ROIs using anatomical masks from a probabilistic atlas (Wang et al., 2015) and combined these for both hemispheres. We included three masks in early visual cortex V1, V2 and V3. V4 and LOC served as ROIs in mid- and high-level ventral visual cortex. We also included four ROIs from dorsal visual cortex: IPS0, IPS1, IPS2 and SPL. We removed all overlapping voxels from these masks to avoid overlap between ROIs. The second step entailed selecting the most activated voxels of the participant-specific *t*-value maps of the localizer run within the previously defined anatomical masks. To keep the number of voxels constant between ROIs and participants, we determined the smallest ROI in any participant when overlaying the localizer *t*-value maps and the anatomical masks. This resulted in a minimum ROI size of 288 voxels. This was then the number of most activated voxels to select of the participant-specific localizer *t*-value maps within all anatomical masks and participants. To select voxels in LOC we used the objects & faces > scrambled contrast and to select voxels in the remaining ROIs we used the objects & scrambled objects > baseline contrast. This resulted in ROI definitions that were specific to each participant with an equal number of voxels across ROIs and participants.

### 5.7 Object location classification from brain measurements

To measure location information in time using EEG and in space using fMRI, we applied multivariate classification (Carlson et al., 2011a; Cichy et al., 2011, 2013; Isik et al., 2014) of object location. Since object location and object category have partly overlapping neural fingerprints in time and space (Cichy et al., 2011; Graumann et al., 2022), we applied a cross-classification scheme that avoided location information results to be confounded with category information (Carlson et al., 2011b; Isik et al., 2014). For this, we cross-classified locations across categories, meaning that during each classification of a given location pair, we trained and tested on different object categories. For all classification analyses described, we employed a binary c-support vector classification (C-SVC) with a linear kernel from the libsvm toolbox (Chang and Lin, 2011) (https://www.csie.ntu.edu.tw/cjlin/libsvm). This cross-classification scheme was applied separately within each background condition, within each attention condition and within each individual participant. The classification scheme was adapted to the specifics of the methods used here: it was applied per time point on the EEG data and per ROI in the fMRI data.

*Time-resolved classification of location from EEG data.* The time-resolved EEG classification analysis (Carlson et al., 2011b; Isik et al., 2014) served to determine the temporal dynamics with which category-independent location information emerged in the brain.

For each time point of the epoched EEG data, we extracted activations from 33 EEG channels. We chose the 33 central and posterior channels starting from the central midline, because we were interested in visual responses and previous studies had shown that location information was most pronounced in those areas (Graumann et al., 2022). We arranged activations from these channels into pattern vectors of 64 conditions and 60 raw trials. Raw trials were randomly arranged into four bins of 15 trials each and averaged by bin into four pseudo-trials to increase SNR. The classification procedure was repeated 100 times, each time assigning random trials into the bins before averaging into pseudo-trials. For classification, three of the pseudo-trials that came from two location conditions of the same category went into the training set. The model resulting from SVM classifier training was then tested on other pseudo-trials coming from the same two location conditions, but from a different category. The accuracy of the classification procedure was measured in percent classification accuracy (50% chance level). This amounted to 6 pairwise location classifications since we had 4 locations that were all classified pairwise once. During each iteration of pairwise location classification, the SVM was trained and tested across all combinations of the four categories in the training and testing set. For example, for a given location classification, the SVM was trained on faces and tested on animals (Fig. 3A). Then the same procedure was applied combing the remaining categories. With four categories in total, this resulted in 6 classification iterations to combine all categories into training and testing pairs. The direction of all training and testing pairs was reversed once (e.g., training on animals and testing on faces and vice versa), yielding a total of 12 classification iterations per pairwise location classification. We averaged 72 (6 location pairs × 12 category train/test pairs) classification accuracies in total per iteration. With 100 iterations with random trial assignment intro pseudo-trials, this resulted in 7,200 classification accuracies that were averaged per background condition, attention condition and participant. The result reflects the amount of location information that is independent of category at each time point, and within a background condition, attention condition and participant.

*Time-resolved EEG searchlight in sensor space*. To gain insights into which EEG channels contained the highest amount of location information we conducted a time-resolved EEG searchlight analysis in EEG channel space. This analysis followed the same scheme as the time-resolved EEG classification described above but extended it by one step: For each EEG channel c, the classification procedure was conducted not on all 33, but on the five closest channels surrounding c. The resulting classification accuracy was stored at the position of c. Iterating across all EEG channels with a temporal resolution downsampled to 10 ms steps, this yielded a map of classification accuracy across all channels and downsampled time points, for each participant, background condition and attention condition.

*Time generalization analysis of location from EEG data*. To characterize the neural dynamics of object location representations across time, we used temporal generalization analysis (Carlson et al., 2011b; Cichy et al., 2014; Isik et al., 2014; King and Dehaene, 2014).

In this analysis, the classification scheme was the same as in the time-resolved EEG classification but with the following extension: besides training and testing the SVM on data from the same time point, we additionally tested the SVM on data from all other time points within a -100 to 600 ms peristimulus time window, downsampled to a 10 ms temporal resolution. This resulted in a two-dimensional matrix of classification accuracies, indexed in rows and columns by the time points of data used for training and testing the SVM. This matrix indicates how much location information was shared at a given combination of time points. This analysis was conducted within time point combination, background condition, attention condition and participant.

*Multivariate fMRI ROI analysis.* The fMRI ROI classification analysis served to determine where category-independent location information emerged in the brain. For each ROI of the fMRI data, we extracted and arranged *t*-values into pattern vectors, one for each of the 16 conditions and 10 runs of the main experiment. Raw trials were randomly arranged into five bins with two runs each and averaged by bin into five pseudo-runs to increase SNR. We then proceeded with a 5-fold leave-one-pseudo-run-out-cross validation procedure. During each classification iteration, we trained an SVM on 4 and tested it on one pseudo-trial. The classification scheme was conceptually equivalent to the EEG classification. Training and testing was conducted across the two different categories, with each being in the training set once. We averaged across the two different training and testing directions of the two categories. The result reflects how much category-tolerant location information was present for each ROI, participant, background and attention condition separately.

### 5.8 Statistical testing

*Wilcoxon signed-rank tests.* To test for above-chance classification accuracy at time points in the EEG time courses, in the EEG time-generalization matrix and for above-chance classification in the fMRI ROI results, we performed non-parametric two-tailed Wilcoxon signed-rank tests. The null hypothesis was always that the parameter being tested (i.e., classification accuracy) came from a distribution with a median of chance level (i.e., 50% classification accuracy for pairwise classification). We corrected the resulting *P*-values for multiple comparisons using false discovery rate at 5% level in every case where more than one test was conducted.

*Bootstrap tests.* To estimate confidence intervals and to compute the significance of peak-to-peak latency differences in the EEG time courses we used bootstrapping. We bootstrapped the participant pool 10,000 times with replacement and calculated the statistic of interest for each of the bootstrap samples.

For the peak-to-peak latency differences in the EEG time courses, we bootstrapped the latency difference between the peaks of the two time courses being compared. This resulted in a bootstrapped distribution that could be compared to zero. To determine the significance of peak-to-peak latencies in the EEG time courses, we computed the proportion of values that were equal to or smaller than zero and corrected them for multiple comparisons using FDR at *P*=0.05. For computing the 95% confidence intervals of peak latencies of each time course, we bootstrapped the peak and computed the 95% percentiles of this distribution.

*ANOVAs.* We used repeated-measures ANOVAs to test for main effects and the interaction between the factors background and attention within ROIs. Since both factors had two levels, the assumption of sphericity was always met.

All post-hoc tests were conducted using pairwise *t*-tests and *P*-values were corrected for multiple comparisons using Tukey correction.

## Data availability

The fMRI and EEG data will be publicly available at the time of publication via https://osf.io/hf6zp/.

## Code availability

Analysis code will be publicly available at the time of publication via https://github.com/graumannm/AttentionLocation.

## Acknowledgements

We thank Benjamin Lahner for the glass brain plots. Computing resources were provided by the high-performance computing facilities at ZEDAT, Freie Universität Berlin. EEG and fMRI data were acquired at the Center for Cognitive Neuroscience, Freie Universität Berlin, Berlin. M.G. and R.M.C. are supported by German Research Council (DFG) (CI241/1-1, CI241/3-1, CI241/7-1). R.M.C. is supported by the European Research Council (ERC-StG-2018-803370). L.A.W. is supported by the University of Konstanz. The funders had no role in study design, data collection and analysis, decision to publish or preparation of the manuscript.

## Author contributions

M.G. and R.M.C. designed research. M.G. and L.A.W. performed experiments. M.G. and L.A.W. performed EEG preprocessing. M.G. performed data analyses. M.G. and R.M.C. wrote the manuscript.

## References

Battistoni E, Kaiser D, Hickey C, Peelen M V (2020) The time course of spatial attention during naturalistic visual search. Cortex 122:225–234.

Boudreau CE, Williford TH, Maunsell JHR (2006) Effects of task difficulty and target likelihood in area V4 of macaque monkeys. J Neurophysiol 96:2377–2387.

Briggs F, Mangun GR, Usrey WM (2013) Attention enhances synaptic efficacy and the signal-to-noise ratio in neural circuits. Nature 499:476–480.

Buffalo EA, Fries P, Landman R, Liang H, Desimone R (2010) A backward progression of attentional effects in the ventral stream. Proc Natl Acad Sci 107:361–365.

Camprodon JA, Zohary E, Brodbeck V, Pascual-Leone A (2010) Two phases of V1 activity for visual recognition of natural images. J Cogn Neurosci 22:1262–1269.

Carlson TA, Hogendoorn H, Fonteijn H, Verstraten FA (2011a) Spatial coding and invariance in object-selective cortex. Cortex 47:14–22.

Carlson TA, Hogendoorn H, Kanai R, Mesik J, Turret J (2011b) High temporal resolution decoding of object position and category. J Vis 11:1–17.

Chang C-C, Lin C-J (2011) Libsvm: A library for support vector machines. ACM Trans Intell Syst Technol 2:1–27.

Chaumon M, Bishop DVM, Busch NA (2015) A practical guide to the selection of independent components of the electroencephalogram for artifact correction. J Neurosci Methods 250:47–63.

Chen Y, Martinez-Conde S, Macknik SL, Bereshpolova Y, Swadlow HA, Alonso JM (2008) Task difficulty modulates the activity of specific neuronal populations in primary visual cortex. Nat Neurosci 11:974–982.

Cichy RM, Chen Y, Haynes JD (2011) Encoding the identity and location of objects in human LOC. Neuroimage 54:2297–2307.

Cichy RM, Pantazis D, Oliva A (2014) Resolving human object recognition in space and time. Nat Neurosci 17:455–462.

Cichy RM, Sterzer P, Heinzle J, Elliott LT, Ramirez F, Haynes J-D (2013) Probing principles of large-scale object representation: Category preference and location encoding. Hum Brain Mapp 34:1636–1651.

Delorme A, Makeig S (2004) EEGLAB: An open source toolbox for analysis of single-trial EEG dynamics including independent component analysis. J Neurosci Methods 134:9– 21.

Desimone R, Duncan J (1995) Selective visual attention. Annu Rev Neurosci 18:193–222.

DiCarlo JJ, Cox DD (2007) Untangling invariant object recognition. Trends Cogn Sci 11:333–341.

Eurich CW, Schwegler H (1997) Coarse coding: Calculation of the resolution achieved by a population of large receptive field neurons. Biol Cybern 76:357–363.

Fahrenfort JJ, Scholte HS, Lamme VAF (2007) Masking disrupts reentrant processing in human visual cortex. J Cogn Neurosci 19:1488–1497.

Gandhi SP, Heeger DJ, Boynton GM (1999) Spatial attention affects brain activity in human primary visual cortex. Proc Natl Acad Sci U S A 96:3314–3319.

Graumann M, Ciuffi C, Dwivedi K, Roig G, Martin R (2022) The spatiotemporal neural dynamics of object location representations in the human brain. Nat Hum Behav:1–38.

Greene M, Oliva A (2009) The Briefest of Glances: The Time Course of Natural Scene Understanding. Psychol Sci 20:464–472.

Groen IIA, Dekker TM, Knapen T, Silson EH (2022) Visuospatial coding as ubiquitous scaffolding for human cognition. Trends Cogn Sci 26:81–96.

Groen IIA, Ghebreab S, Lamme VAF, Scholte HS (2016) The time course of natural scene perception with reduced attention. J Neurophysiol.

Groen IIA, Jahfari S, Seijdel N, Ghebreab S, Lamme VAF, Scholte HS (2018) Scene complexity modulates degree of feedback activity during object detection in natural scenes. PLoS Comput Biol 14:e1006690.

Guggenmos M, Sterzer P, Cichy RM (2018) Multivariate pattern analysis for MEG: A comparison of dissimilarity measures. Neuroimage 173:434–447.

Herrero JL, Gieselmann MA, Sanayei M, Thiele A (2013) Attention-induced variance and noise correlation reduction in macaque V1 is mediated by NMDA receptors. Neuron 78:729–739.

Hillyard SA, Teder-Sälejärvi WA, Münte TF (1998a) Temporal dynamics of early perceptual processing. Curr Opin Neurobiol 8:202–210.

Hillyard SA, Vogel EK, Luck SJ (1998b) Sensory gain control (amplification) as a mechanism of selective attention: Electrophysiological and neuroimaging evidence. Philos Trans R Soc B Biol Sci 353:1257–1270.

Hong H, Yamins DLK, Majaj NJ, DiCarlo JJ (2016) Explicit information for category-orthogonal object properties increases along the ventral stream. Nat Neurosci 19:613– 622.

Isik L, Meyers EM, Leibo JZ, Poggio T (2014) The dynamics of invariant object recognition in the human visual system. J Neurophysiol 111:91–102.

Itthipuripat S, Sprague TC, Serences JT (2019) Functional MRI and EEG index complementary attentional modulations. J Neurosci 39:6162–6179.

Kaiser D, Oosterhof NN, Peelen M V. (2016) The neural dynamics of attentional selection in natural scenes. J Neurosci 36:10522–10528.

Kaiser D, Quek GL, Cichy RM, Peelen M V. (2019) Object vision in a structured world. Trends Cogn Sci 23:672–685

Kar K, Kubilius J, Schmidt K, Issa EB, DiCarlo JJ (2019) Evidence that recurrent circuits are critical to the ventral stream’s execution of core object recognition behavior. Nat Neurosci 22:974–983.

Kay KN, Weiner KS, Grill-Spector K (2015) Attention reduces spatial uncertainty in human ventral temporal cortex. Curr Biol 25:595–600.

Khayat PS, Spekreijse H, Roelfsema PR (2006) Attention lights up new object representations before the old ones fade away. J Neurosci 26:138–142.

King JR, Dehaene S (2014) Characterizing the dynamics of mental representations: The temporal generalization method. Trends Cogn Sci 18:203–210.

Koivisto M, Railo H, Revonsuo A, Vanni S, Salminen-Vaparanta N (2011) Recurrent processing in V1/V2 contributes to categorization of natural scenes. J Neurosci 31:2488–2492.

Kravitz DJ, Saleem KS, Baker CI, Mishkin M (2011) A new neural framework for visuospatial processing. Nat Rev Neurosci 12:217–230.

Lakatos P, Karmos G, Mehta AD, Ulbert I, Schroeder CE (2008) Entrainment of neuronal oscillations as a mechanism of attentional selection. Science 320:110–113.

Lamme VAF, Roelfsema PR (2000) The distinct modes of vision offered by feedforward and recurrent processing. Trends Neurosci 23:571–579.

Lavie N (1995) Perceptual Load as a Necessary Condition for Selective Attention. J Exp Psychol Hum Percept Perform 21:451–468.

Lee J, Maunsell JHR (2010) Attentional modulation of MT neurons with single or multiple stimuli in their receptive fields. J Neurosci 30:3058–3066.

Luck SJ, Woodman GF, Vogel EK (2000) Event-related potential studies of attention. Trends Cogn Sci 4:432–440.

Mangun GR (1995) Neural mechanisms of visual selective attention. Psychophysiology 32:4–18.

Martínez A, DiRusso F, Anllo-Vento L, Sereno MI, Buxton RB, Hillyard SA (2001) Putting spatial attention on the map: Timing and localization of stimulus selection processes in striate and extrastriate visual areas. Vision Res 41:1437–1457.

Milner AD, Goodale MA (2006) The visual brain in action. Oxford: Oxford University Press.

Murray SO, Wojciulik E (2004) Attention increases neural selectivity in the human lateral occipital complex. Nat Neurosci 7:70–74.

Noesselt T, Hillyard SA, Woldorff MG, Schoenfeld A, Hagner T, Jäncke L, Tempelmann C, Hinrichs H, Heinze HJ (2002) Delayed striate cortical activation during spatial attention. Neuron 35:575–587.

Oliva A (2005) Gist of the scene. Elsevier Inc.

Peelen M V., Kastner S (2011) A neural basis for real-world visual search in human occipitotemporal cortex. Proc Natl Acad Sci U S A 108:12125–12130.

Peelen M V., Kastner S (2014) Attention in the real world: Toward understanding its neural basis. Trends Cogn Sci 18:242–250

Rajaei K, Mohsenzadeh Y, Ebrahimpour R, Khaligh-Razavi S-M (2019) Beyond core object recognition: Recurrent processes account for object recognition under occlusion. PLOS Comput Biol 15:e1007001.

Reddy L, Kanwisher N (2007) Category Selectivity in the Ventral Visual Pathway Confers Robustness to Clutter and Diverted Attention. Curr Biol 17:2067–2072.

Reynolds JH, Chelazzi L (2004) Attentional modulation of visual processing. Annu Rev Neurosci 27:611–647.

Roelfsema PR, Lamme VAF, Spekreijse H (1998) Object-based attention in the primary visual cortex of the macaque monkey. Nature 395:376–381.

Seijdel N, Loke J, van de Klundert R, van der Meer M, Quispel E, van Gaal S, de Haan EHF, Scholte HS (2021) On the necessity of recurrent processing during object recognition: It depends on the need for scene segmentation. J Neurosci 41:6281–6289.

Seijdel N, Tsakmakidis N, De Haan EHF, Bohte SM, Scholte HS (2020) Depth in convolutional neural networks solves scene segmentation. PLoS Comput Biol 16:e1008022

Silver MA, Ress D, Heeger DJ (2005) Topographic maps of visual spatial attention in human parietal cortex. J Neurophysiol 94:1358–1371.

Spitzer H, Desimone R, Moran J (1988) Increased attention enhances both behavioral and neuronal performance. Science 240:338–340.

Spoerer CJ, Kietzmann TC, Mehrer J, Charest I, Kriegeskorte N (2020) Recurrent neural networks can explain flexible trading of speed and accuracy in biological vision. PLoS Comput Biol 16:e1008215.

Sprague TC, Serences JT (2013) Attention modulates spatial priority maps in the human occipital, parietal and frontal cortices. Nat Neurosci 16:1879–1887.

Szczepanski SM, Konen CS, Kastner S (2010) Mechanisms of spatial attention control in frontal and parietal cortex. J Neurosci 30:148–160.

Tang H, Buia C, Madhavan R, Crone NE, Madsen JR, Anderson WS, Kreiman G (2014) Spatiotemporal dynamics underlying object completion in human ventral visual cortex. Neuron 83:736–748.

Tang H, Schrimpf M, Lotter W, Moerman C, Paredes A, Caro JO, Hardesty W, Cox D, Kreiman G (2018) Recurrent computations for visual pattern completion. Proc Natl Acad Sci U S A 115:8835–8840.

Treisman AM, Gelade G (1980) A feature-integration theory of attention. Cogn Psychol 12:97–136.

Ungerleider L, Haxby J V. (1994) “What” and “where” in the human brain. Curr Opin Neurobiol 4:157–165.

van Bergen RS, Kriegeskorte N (2020) Going in circles is the way forward: the role of recurrence in visual inference. Curr Opin Neurobiol 65:176–193.

VanRullen R, Thorpe SJ (2001) The time course of visual processing: From early perception to decision-making. J Cogn Neurosci 13:454–461.

Võ MLH, Boettcher SE, Draschkow D (2019) Reading scenes: How scene grammar guides attention and aids perception in real-world environments. Curr Opin Psychol 29:205– 210.

Wandell BA, Winawer J (2015) Computational neuroimaging and population receptive fields. Trends Cogn Sci 19:349–357.

Wang L, Mruczek REB, Arcaro MJ, Kastner S (2015) Probabilistic maps of visual topography in human cortex. Cereb Cortex 25:3911–3931.

Wolfe JM (1994) Visual search in continuous, naturalistic stimuli. Vision Res 34:1187–1195.

Wolfe JM, Võ MLH, Evans KK, Greene MR (2011) Visual search in scenes involves selective and nonselective pathways. Trends Cogn Sci 15:77–84.

Wyatte D, Jilk DJ, O’Reilly RC (2014) Early recurrent feedback facilitates visual object recognition under challenging conditions. Front Psychol 5:1–10.

