## Supplementary Material for "Independent spatiotemporal effects of spatial attention and background clutter on human object location representations"

#### Supplementary Figure 1: Time generalization analysis scheme and results for classifying object location across time, category and background condition

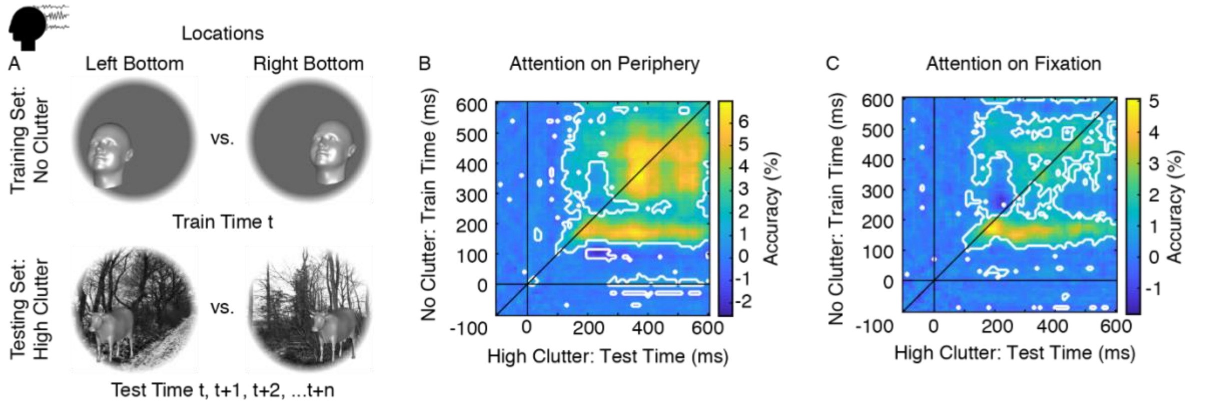


**Supplementary Figure 1. A,** Time generalization analysis scheme for classifying object location across time, category and background condition. The classification scheme was the same as in Fig. 3A with the difference that training set conditions always came from the no clutter and testing set conditions came from the high clutter condition, within attention condition. Objects are enlarged for visibility and did not extend into another quadrant in the original stimuli. **B,** Location classification across categories, background conditions and time points in the peripheral attention condition. Horizontal and vertical black lines indicate stimulus onset, the oblique black line highlights the diagonal. White outlines indicate significant time points ($N$=26, two-tailed Wilcoxon signed-rank test, $P$<0.05, FDR corrected). The classification peaks were significantly shifted below the diagonal by 69 ms ($N$=26, two-sided Wilcoxon-signed-rank test, $P$<0.001), indicating that an early process in the no clutter condition generalized to later time points in the high clutter condition (Supplementary Methods 1). **C,** Location classification across categories, background conditions and time points in the fixation attention condition. The classification peaks were also significantly shifted below the diagonal by 35 ms in the fixation attention condition ($N$=26, two-sided Wilcoxon-signed-rank test, $P$=0.03), indicating a shared, delayed processing stage between location representations in the no and high clutter conditions (Supplementary Methods 1).

### Supplementary Methods 1: Computation of peak-to-diagonal distance in the time-generalization matrices

To statistically test whether the peak in classification accuracy was significantly higher above or below the diagonal in the time-generalization matrices across background conditions (Supplementary Fig. 1B,C), we computed the distance between the classification peak and the diagonal for each participant. To compute the distance between the diagonal and classification peak in the time-generalization matrix, we first determined the peak coordinates $(p_x, p_y)$ along the x- and y-axes in the post-stimulus time window. Then we calculated the coordinates of the point on the diagonal that was closest to the peak by calculating $b_x$:

$$b_x = \frac{(p_x + p_y)}{2}$$

This gave the coordinates for the closest point on the diagonal since on the diagonal, x and y coordinates are the same. With these coordinates we calculated the shortest perpendicular Euclidean distance between the peak and the closest point on the diagonal:

$$d_{euclidean} = \sqrt{(p_x - b_x)^2 + (p_y - b_x)^2}$$

Since Euclidean distances are always positive and we tested the computed distances for a bias towards upper or lower diagonal against chance level, we set

$$d_{euclidean} = d_{euclidean} \times -1$$

for all cases where $p_x < p_y$ which is the case for all peaks above the diagonal. This allowed us to statistically test Euclidean distances against 0.
